# Repetitive neonatal pain increases spinal cord DNA methylation of the µ-opioid receptor

**DOI:** 10.1101/2024.08.25.609598

**Authors:** Mathilde Baudat, Elbert A.J Joosten, Sinno H.P. Simons, Daniël L.A. van den Hove, Renzo J.M. Riemens

## Abstract

Repetitive neonatal painful procedures experienced in the neonatal intensive care unit (NICU) are known to alter the development of the nociceptive system and have long-lasting consequences, notably lower post-operative µ-opioid receptor levels in the spinal cord. Given the influence of the NICU on the epigenome, the present study hypothesized that neonatal procedural pain alters the DNA methylation status of the opioid receptor mu 1 encoding gene (*Mor-1*) in the spinal cord and dorsal root ganglions (DRGs). To this end, the needle prick model of repetitive neonatal pain was used, and methylation of *Mor-1* promotor was assessed in the spinal cord and the DRG using bisulfite pyrosequencing. Our findings demonstrated that neonatal procedural pain increased spinal cord *Mor-1* promotor DNA methylation in the ipsilateral side as compared to the contralateral side, an effect that was not observed in the control animals, nor in the DRG. We also identified a behaviorally-associated CpG site following neonatal needle pricks. This study is the first to highlight a localized and noxious-stimuli-dependent effect of repetitive neonatal procedural pain on *Mor-1* promotor methylation and emphasizes the need to explore the effects of repetitive neonatal procedural pain on the epigenome.

**Impact:**

- This study reveals that repetitive neonatal procedural pain increases DNA methylation of the Mor-1 promoter in the spinal cord of neonatal rats.
- This is the first study to identify an effect of neonatal procedural pain on DNA methylation, emphasizing the critical need for further investigation into the epigenetic consequences of neonatal procedural pain.
- These insights could lead to better management and treatment strategies to mitigate the long-term impacts of early pain exposure on neurodevelopment and behavior.

## Introduction

Neonatal pain is a topic of raising importance in light of the steady increase in premature birth (before 37 weeks of gestation) and neonatal intensive care until (NICU) admission incidences, (1–3). The 10% of newborns admitted to the NICU undergo an average of 10-14 daily painful procedures (3–5). This repetitive stimulation of nociceptive circuits by painful procedures takes place at a time of neurodevelopmental vulnerability and maturation and leads to inadequate programming of the pain network, including the descending inhibitory system (6–9).

The opioid system is pivotal in controlling the local spinal nociceptive circuits and is still maturing during the neonatal period (10,11). The most important opioid receptor in the descending control of nociception is the µ-opioid receptor 1 (MOR-1), localized on the pre-and post-synaptic membrane of primary and secondary afferent nociceptive fibers, respectively localized in the dorsal root ganglion (DRG) and spinal cord (12,13). Neonatal pain is not without effects on the descending opioid pathways: a single neonatal inflammatory event on the day of birth was shown to increase opioid tone and decrease MOR-1 expression in the periaqueductal grey (PAG) of adult rats (9). Moreover, previous work by our group has shown that adult animals previously exposed to neonatal procedural pain developed decreased protein levels of MOR-1 in the spinal dorsal horn following adult injury, without changes in baseline MOR-1 (14). These results suggest that repetitive neonatal procedural pain may prime the endogenous opioid system in the spinal dorsal horn, which becomes apparent only after a second injury in the adult.

One plausible underlying mechanism is the alteration of the epigenome. Epigenetics encompasses the mechanisms able to regulate gene transcription without altering the primary DNA sequence. To this day, the most extensively studied epigenetic process is DNA methylation which occurs at cytosine-phosphate-guanine sites (CpGs). Methylation of CpGs may regulate gene expression by orchestrating the binding of transcription factors or by locally inducing changes to the chromatin structure, thereby affecting its accessibility and, as such, transcriptional activity and protein expression (15). Upon early life adversity, the epigenome may be reprogrammed to adapt to the environment, which could result in neurodevelopmental alterations that predispose or prime individuals to later-life diseases (16,17).

Recently, clinical studies revealed that NICU stay significantly decreases methylation levels of *Mor-1*, while increasing methylation of the serotonin transporter gene (*SLC6A4*) (18–20). Furthermore, *Mor-1* was shown to be sensitive to epigenetic regulation after neuropathic nerve injury in adult mice (21). Given these findings, we hypothesized that repetitive neonatal procedural pain alters the methylation of the *Mor-1* promotor in the DRG and the spinal cord. In order to test this hypothesis, we quantified the methylation levels of the *Mor-1* promotor in the spinal cord and the DRG of neonatal rats exposed to repetitive neonatal procedural pain, using bisulfite-pyrosequencing.

## Materials and methods

### Ethics statement

All animal experiments were performed in accordance with the European Directive for Protection of Vertebrate Animal Use for Experimental and Other Scientific Purposes (86/609/EEC) and were approved by the Committee for Experiments on Animals, Maastricht, The Netherlands (DEC 2017-017).

### Animals

For this study, 32 Sprague Dawley (SD) male and female rat pups from four time-pregnant SD dams were used (Charles River). Breeding of dams was performed at the animal facilities of Maastricht University. On the day of birth, referred to as post-natal day 0 (P0), the litters were culled to a maximum of N=10, and pups were randomly assigned to neonatal conditions using an online randomization tool (random.org; Table 1). All animals were housed in a room with controlled temperature (19-24 °C) and humidity (55 ± 15%), a reversed 12h/12h day-night cycle, and background music. *Ad libitum* water and food were available throughout the whole study period.

**Table 1.** Animal and sample distribution. Litters were either assigned to the undisturbed control (UC) group, or to the treatment litter. Pups in the treatment litter were randomly assigned on the day of birth to either the needle prick (NP) or disturbed control (DC) group. At P10 animals were sacrifice and lumbar spinal cord and L4-L6 DRG were dissected. 2 samples were excluded from the *Mor-1* promotor methylation analysis due to low pyrosequencing signal. Ipsi.: ipsilateral; Contra.: contralateral.

|  | Behavior |  |  | Average *Mor-1* promotor methylation |  |  |  |
| --- | --- | --- | --- | --- | --- | --- | --- |
|  |  |  |  | Spinal cord |  | DRG |  |
|  | Male | Female | Total | Ipsi. | Contra. | Ipsi. | Contra. |
| NP | 6 | 4 | 10 | 10 | 10 | 10 | 10 |
| DC | 7 | 4 | 11 | 11 | 11 | 10 | 10 |
| UC | 4 | 7 | 11 | 11 | 10 | 10 | 9 |

### Neonatal procedures

To model repetitive procedural pain exposure in the NICU, a repetitive neonatal needle prick model was implemented as previously described by Knaepen and colleagues (22). Newborn pups were noxiously stimulated four times a day via unilateral 2mm calibrated needle pricks in the mid-plantar surface of the left hind paw from P0 to P7 (needle prick, NP, N=10; Figure 1). Control animals were shortly handled at the same hourly intervals as the NP animals (disturbed control, DC, N=11) or were left undisturbed (undisturbed control, UC, N=11).

**Figure 1.**
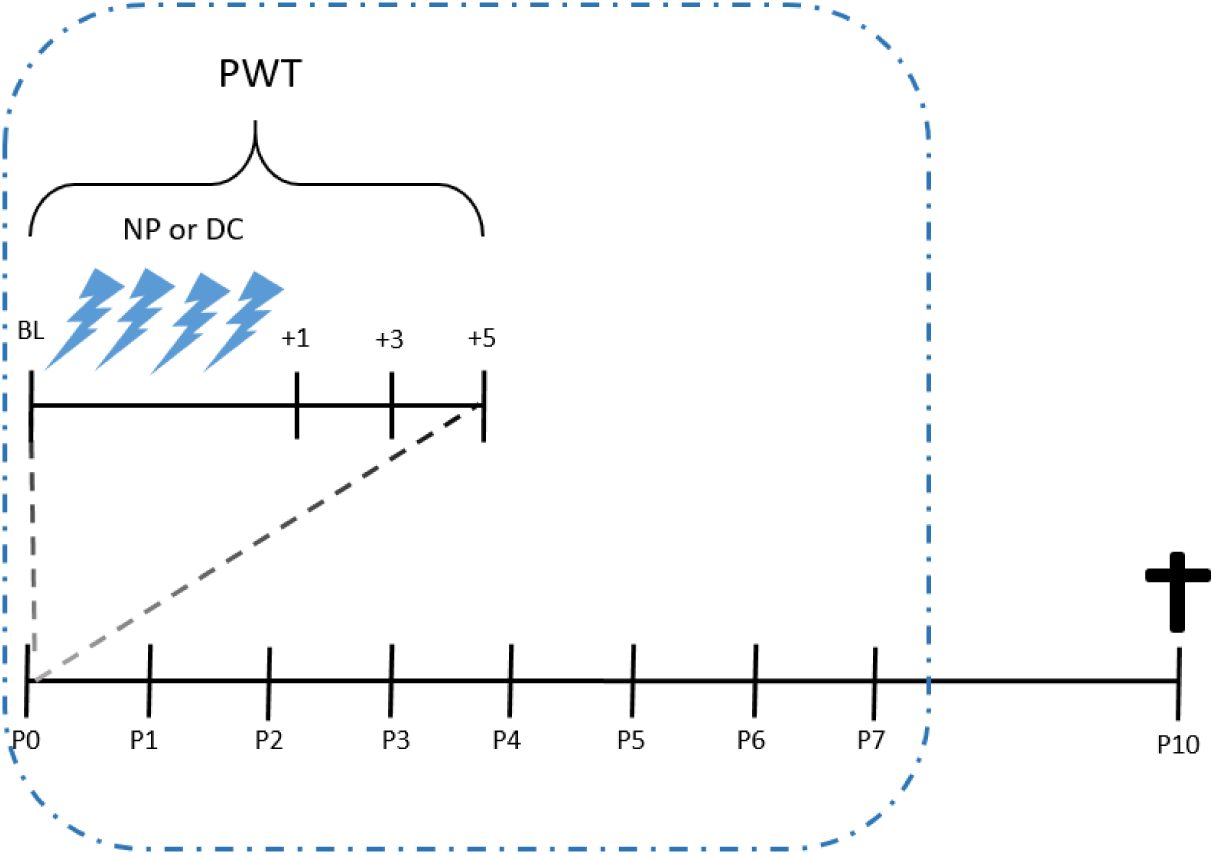
Experimental design and timeline. From postnatal day 0 (P0) to P7, animals were either needle pricked (NP, N=10), handled (DC, N=11) 4 times a day or were left undisturbed (UC, N=11). Paw withdrawal threshold (PWT) was assessed before (Baseline; BL) and 1, 3 and 5 hours (+1, +3, +5) after the last needle prick or handling using von Frey filaments. At P10 animals were sacrificed and spinal cord and DRGs were dissected.

### Von Frey for mechanical sensitivity in neonates

Paw withdrawal thresholds (PWT) of the ipsi- and contralateral hind-paws were assessed before (Baseline; BL) and 1, 3, and 5 hours after the last noxious stimulation or handling using dorsal von Frey. Ascending Von Frey filaments (bending force 0.407g, 0.692g, 1.202g, 2.041g, 3.63g from P4 onwards and 5.495g from P6 onwards; Stoelting, Wood Dale, IL, USA) were applied 5 times to the dorsal surface of the hind-paws. The number of positive responses, that is paw withdrawal or flinching behavior evoked by the filaments, per filament, was recorded, and behavioral testing was discontinued when five positive responses were observed. A 50% PWT was calculated using a sigmoidal curve fitting in GraphPad Prism 9.5.1 (GraphPad Software, San Diego, USA).

### Tissue collection

At P10, all animals (N=32) were weighted and randomly and alternatively collected from each nest and sacrificed. Animals were decapitated and dissected to collect the lumbar spinal cord (N=32) and the DRG (N=30) followed by snap-freezing in liquid nitrogen. Only the lumbar part of the spinal cord was collected and ipsi- and contra-lateral sides were separated. Ipsi- and contra-lateral L4 to L6 DRG were collected and pooled. All tissues were stored at −80℃ until further processing.

### DNA isolation and purification

DNA was extracted using a standard Phenol/Chloroform-Isoamyl alcohol (PCI) extraction method. In brief, the samples were lysated in 500 µl lysis buffer containing 50 mM Tris (pH 8.0), 1 mM EDTA and 0.5% SDS. After adding 25 µl of proteinase K (Thermo Fisher Scientific, Waltham, MA, USA), the samples were incubated overnight at 56°C in a shaking thermoblock. Following incubation, the proteinase K was inactivated at 80°C for 10 minutes. PCI (#77617-100, Sigma Aldrich, Saint Louis, MO, USA) was added in a 1:1 ratio, the samples were manually mixed for 5 minutes and then centrifuged at 14.000 rpm for 5 minutes. The upper phase was carefully transferred to a new sterile 1.5ml Eppendorf tube. Another equal volume (1:1 ratio) of PCI was added, after which the samples were mixed for 5 minutes and centrifuged at 14.000 rpm for 5 minutes. The upper phase from each sample was once again transferred to a new sterile 1.5 ml Eppendorf tube. The DNA was precipitated by adding 50µl of 3 M NaAc (pH 5.6) and 1250µl of 100% cold (−20°C) ethanol, then incubated for at least 30 minutes at - 80°C and centrifuged for 30 minutes at 14.000 rpm at 4°C. Subsequently, the solution was carefully removed, and the DNA pellets were washed using 70% cold ethanol and centrifuged for 5 minutes at 14.000 rpm at 4°C. Next, the ethanol was carefully removed and the DNA pellets were air-dried at room temperature. Finally, the isolated DNA from each sample was dissolved in a volume of 50µL Milli-Q and then stored at −20°C until further processing. The DNA yield of each sample was then quantified by using a Qubit dsDNA HS Assay Kit (Invitrogen, Waltham, MA, USA).

### DNA bisulfite conversion

The EZ DNA Methylation-Gold Kit (#D5008, Zymo Research, Irvine, CA, USA) was used according to the manufacturer’s instructions to bisulfite convert 400ng of each DNA sample. Briefly, DNA was incubated with the CT conversion reagent for 8 minutes at 98°C followed by a 150-minute incubation at 64°C and a final storage step at 4°C. After the bisulfite clean-up procedure, each sample was collected in a single 1.5 mL Eppendorf tube by flushing the spin column twice using 20 µL of elution buffer, resulting in a final concentration of 10 ng/µL for each fraction when assuming full recovery of the bisulfite-converted DNA. Samples were randomized across bisulfite conversion plates and processed simultaneously to avoid batch effects. Bisulfite-converted DNA was aliquoted and stored at −20°C until further processing.

### Polymerase chain reaction

Primers targeting the *Mor-1* promotor were designed using the PyroMark Assay Design 2.0 software (Qiagen, Hilden, Germany) and were based on the Ensembl mRatBN7.2 genome build (Table S1). Two assays were developed to increase the coverage of the targeted region. Polymerase chain reaction (PCR) amplification of the targeted region was performed with an initial denaturation step at 95°C for 5 minutes, followed by 55 cycles at 95°C, 56°C and 72°C for 30, 30, and 30 seconds, respectively, with a final extension step at 72°C for 1 minute. For each individual PCR reaction, 10ng of the bisulfite-converted DNA was used. Each of the reactions contained 2.5 μL PCR buffer (10X) with 20 mM MgCl2, 0.5 μL 10 mM dNTP mix, 1 μL of each respective primer (5 μM stock) and 0.2 μL (5 U/μL) FastStart Taq DNA Polymerase (Roche Diagnostics GmbH, Mannheim, Germany) in a total volume of 25 μL. The PCR products were visualized on a 2% agarose gel and 10 μL product was utilized per assay for bisulfite pyrosequencing.

### Pyrosequencing

The Pyromark Q48 Autoprep system with the PyroMark Q48 Advanced CpG Reagents (Qiagen) and PyroMark Q48 Magnetic Beads (Qiagen) were used for bisulfite pyrosequencing according to the manufacturer’s instructions. All pyrosequencing assays were tested for their sensitivity on various fractions, *i.e.*, 0%, 25%, 50%, 75%, and 100% of methylated DNA standards that were generated from the rat premixed calibration standard set (80-8060R-PreMx, EpigenDx, Hopkinton, MA, USA). Modification levels at a single CpG resolution were analyzed with the Pyromark Q48 Autoprep software (Qiagen).

### Statistics

Differences in mechanical sensitivity during the neonatal period were analyzed using a repeated measure analysis of variance (ANOVA; assessing the effects of age and condition) with Holm-Sidak post-hoc correction. The difference in mechanical sensitivity was analyzed using GraphPad Prism 9.5.1 (GraphPad Software, San Diego, USA) and results were considered significant at p<0.05.

Methylation levels were analyzed using a mixed ANOVA model. Neonatal condition was defined as a between-subject variable, and side (ipsi- vs contra-lateral) was considered a within-subject variable. When a significant effect for the between- or the within-subject variable was obtained, subsequent t-tests were performed for relevant comparisons only. Spinal cord and DRG data were analyzed separately. Average methylation levels over the two assays (assay 1: Ensembl, mRatBN7.2, 1:49708770-49708956; assay 2: Ensembl, mRatBN7.2, 1:49708984-49709078), were calculated by averaging the methylation levels of all CpGs per animal. Following bisulfite pyrosequencing, two samples did not meet the quality control standards of the Pyromark Q48 Autoprep software and were therefore excluded from the analysis (Table 1). Analysis of methylation differences for each CpG was done using a repeated measure mixed-ANOVA, with between and within-subject variables as described above, and CpGs as repeated measures. To explore if behavioral data was correlating with *Mor-1* spinal methylation level, a Spearman correlation analysis was performed between average spinal *Mor-1* promotor methylation and CpG sites methylation levels. The behavioral data used for the correlation analysis was the last PWT, that is at P7, 5 hours after the last needle prick or short maternal separation. The mixed ANOVA and Spearman correlation were analyzed using SPSS version 28.0.1.1. and results were considered significant at p<0.05. All data are presented as mean ± standard error of the mean (SEM), and created with GraphPad Prism 9.5.1.

## Results

### Neonatal needle pricks, but not handling, decrease the ipsilateral mechanical paw withdrawal threshold

From P0 to P7, all 32 animals were included in the study and subjected to either needle pricks (NP, N=10), or handling (DC, N=11), or were left undisturbed (UC, N=11). To verify the model (22–25), mechanical sensitivity was tested daily by von Frey tests before 1, 3, and 5 hours after the last needle pricking or handling session. As sex did not significantly affect the ipsilateral (F_1,17_=0.1327, p=0.72) or contralateral (F_1,17_=3.904, p=0.064) baseline PWT at P0, and no-sex effect were previously notes in the needle prick model (14,23,25), males and females were pooled to increase power. Ipsilateral PWT was shown to significantly increase over time (F_31, 589_=23.56, p<0.001; Figure 2), as expected with thickened skin texture and increasing weight (F_1.930, 41.62_=612.8, p<0.001). The latter was not affected by the neonatal condition (F_2,29_=2.813, p=0.076). Neonatal condition significantly affected ipsilateral PWT (F_1, 19_=13.29, p=0.002), and an interaction effect between time and neonatal condition was observed (F_31, 589_=8.371, p<0.001). Post-hoc analysis revealed that NP animals had significantly lower PWT from P5+5h onwards when compared to DC animals. On the contralateral side, only age affected the PWT (F_31,589_=50.88, p<0.0001).

**Figure 2.**
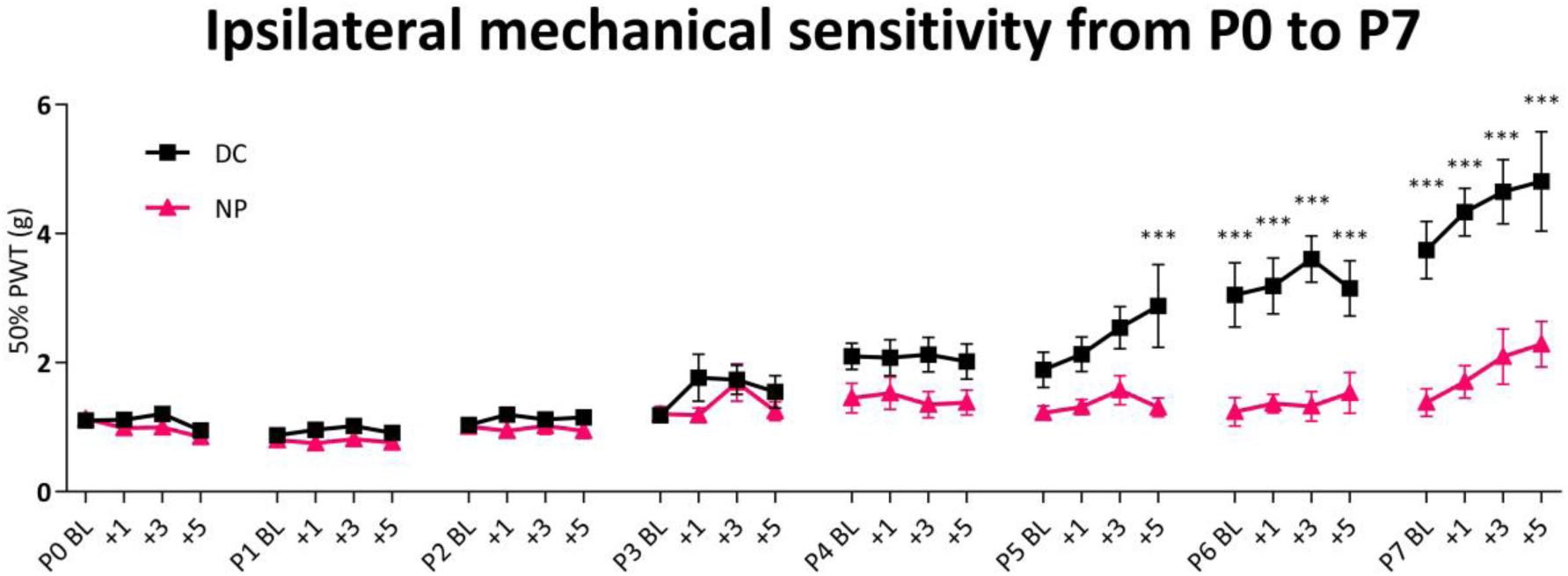
Mechanical sensitivity after needle prick or handling during the neonatal week. From post-natal day 5 (P5), repetitive needle prick (NP, N=10) results in decreased ipsilateral paw withdrawal threshold (PWT) compared to handling (DC, N=11; F_1, 19_=13.29, p=0.002). From P0 to P7, PWT significantly increased over time independently of neonatal condition (F_31,589_=26.56, p<0.001). Postnatal day 0 to 7 (P0-7); BL, baseline von Frey measurement; +1/+3/+5, von Frey measurement 1/3/5 hours after last needle pricks on P0-7. Data plotted as mean ± SEM. *** p<0.001.

### Neonatal needle pricking increases methylation of the Mor-1 promotor in the spinal cord, but not the DRG

Following sacrifice (N=32), the ipsilateral and contralateral spinal cord (N=32) and DRG of P10 animals (N=30) were dissected and DNA was extracted for bisulfite pyrosequencing of the *Mor-1* promotor. Side (ipsilateral vs contralateral) significantly affected the methylation of *Mor-1* promoter in the spinal cord (F_1,28_=5.690, p=0.024), but not the DRG (F_1,26_=0.709, p=0.408). A paired t-test revealed that this laterality effect was only observed in animals that underwent neonatal needle pricks from P0 to P7 (t9=2.683, p=0.025; Figure 3). No interaction effect between neonatal condition and side was observed in the spinal cord (F_2,28_=1.02, p=0.374) or the DRG (F_2,26_=0.292, p=0.749) methylation levels of the *Mor-1* promotor.

**Figure 3.**
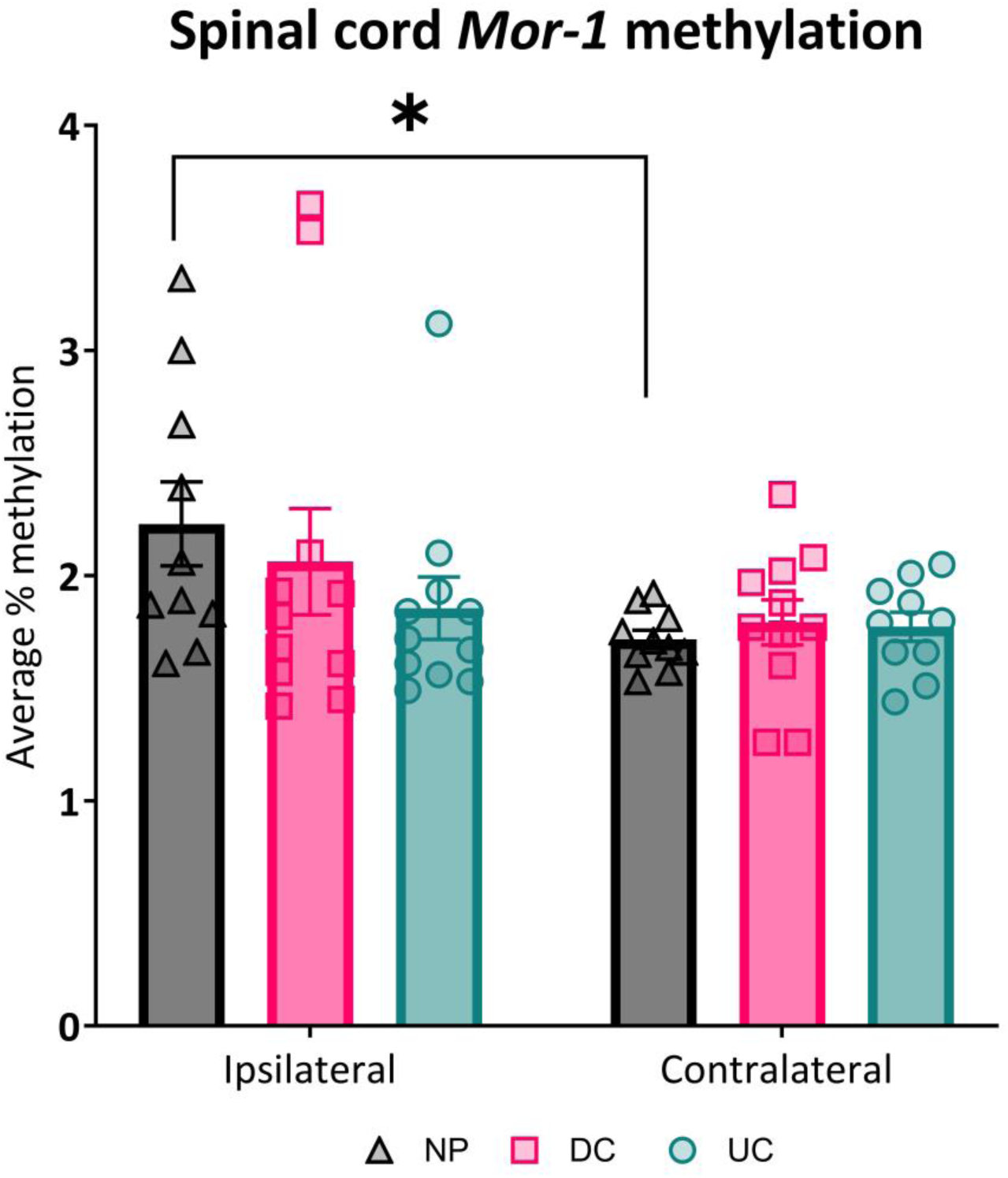
Spinal cord methylation levels at P10 of the *Mor-1* promotor. At P10, the *Mor-1* promotor was significantly hyper-methylated by neonatal needle pricks (NP, N=10) on the ipsilateral side, but not on the contralateral side (t9=2.683, p=0.025). This effect was not observed in the DC (N= 11) or UC animals (N=11). Data plotted as mean ± SEM. *p=0.025.

In order to identify which region of the promotor was affected by needle pricks, a mixed model ANOVA was performed on the average methylation per assay, and for each CpG site. In the spinal cord, the effect of neonatal needle pricking on the methylation of *Mor-1* promotor was localized between 1:49708984 and 1:49709078 (F_1,28_=5.291, p=0.029). Methylation levels were dependent on the CpG sites in the spinal cord (F_8,224_=12.949, p=0.00) and the DRG (F_8,208_=62.827, p=0.00), and no CpG site showed a significant difference between the ipsilateral and the contralateral side (Table S2). The Spearman correlation analysis revealed that methylation of the CpG 4, located at 1:49709023-49709024, was significantly correlated with the ipsilateral PWT in the NP animals (R_s_ =0.855, p=0.002, Table S4). No other significant correlations were observed (Table S4). Neither neonatal condition (F_2,26_=1.965, p=0.106) nor side (F_1,26_=0.709, p=0.408) affected methylation levels in the DRG (Table S3).

## Discussion

The present study aimed to investigate the effects of repetitive neonatal procedural pain on methylation of the *Mor-1* promotor in the rat spinal cord and the DRG. Our findings highlight the successful induction of repetitive neonatal procedural pain and reveal that repetitive needle pricks increased *Mor-1* promotor methylation in the ipsilateral spinal cord at P10. This effect was confined to a specific region of the *Mor-1* promotor (1:49708984-49709078). The absence of methylation changes at the *Mor-1* promotor region in the DRG implies a localized impact of repetitive noxious stimulation on the developing nociceptive system. The DRG contain nuclei of primary afferent fibers, where activation of pre-synaptic MOR-1 inhibits excitatory transmission (12,13,26). On the other hand, DNA collected from the spinal cord originates from nuclei of second-order nociceptive neurons, where MOR-1 is localized post-synaptically and leads to hyperpolarization of nociceptive neurons sending noxious signals to the brain (12). The effect of repetitive stimulation of the nociceptive circuit in the spinal cord, but not the DRG, indicates a localized epigenetic regulation of post-synaptic MOR-1 by repetitive neonatal procedural pain.

In recent years, clinical studies started to investigate the consequences of NICU admission on the epigenome (18–20,27). Two of those studies investigated *MOR-1* and reported conflicting results: one described hypo-methylation of *MOR-1* upon NICU discharge (18), while another found no effect (27). In both reports, DNA methylation was measured using systemic samples (stool and saliva, respectively) and therefore does not reflect epigenetic changes on local nociceptive pathways. Furthermore, conclusions concerned the stressful NICU environment, and therefore do not provide evidence of the effects related to neonatal procedural pain. The inherent constraints of clinical studies like these, despite their indisputable value, highlight the imperative of employing preclinical models to further elucidate the consequences of neonatal procedural pain on the developing nociceptive system. In line with Hatfield and colleagues (27), the present study provides evidence that methylation of the *Mor-1* promotor is noxious-stimulus-dependent and is not affected by the neonatal stressful environment (e.g. handling, repetitive maternal separation, maternal stress), as shown by the lack of differences between disturbed and undisturbed animals.

The enriched methylation of the *Mor-1* promotor in response to repetitive neonatal procedural pain ties in with alterations within the descending inhibitory opioid pathway following neonatal pain. An enhanced opioid tone in the PAG was observed in rats exposed to neonatal inflammatory pain and drives the ensuing long-term hypo-algesia (9). Furthermore, MOR-1 expression and binding within the PAG were diminished following the rapid internalization of the receptor by endogenous opioid activation (9). Although LaPrairie and colleagues investigated neonatal inflammatory pain, decreased MOR-1 expression in the spinal cord was also observed after adult injury in a rat model of repetitive neonatal procedural pain (14). Previous studies have shown that methylation of the *Mor-1* promotor correlates with downregulation of the MOR-1 gene (28,29). Following repetitive stimulation of the developing nociceptive circuits, methylation of the *Mor-1* promotor displayed a small but significant increase. We previously reported that animals exposed to repetitive neonatal procedural pain reduced the intensity of MOR-1-immunoreactivity in the spinal cord after adult surgery (14). This effect was observed in both the ipsilateral and contralateral spinal cord, indicating that the hyper-methylation as observed in our study could play a role in MOR-1 expression regulation. Nevertheless, other mechanisms are likely also involved in the regulation of MOR-1 after repetitive neonatal procedural pain. Distorted regulation of MOR-1 by cytokines is plausible given that pain experienced after neonatal surgery primes spinal microglia, which are the main source of cytokines in the spinal cord (30). Taken together, our current hypothesis is that repetitive noxious stimulation of the developing nociceptive circuits enhances endogenous opioid tones. The resulting hyper-methylation of the *Mor-1* promotor would then prompt, along with other regulatory mechanisms, the depletion of MOR-1 following adult injury.

Alterations in the epigenome during the development of the nociceptive network are likely part of broader developmental programming induced by repetitive neonatal procedures. For instance, clinical studies have long shown that exposure to painful interventions in neonates results in altered nociceptive processing in adolescents and young adults (31,32). Furthermore, methylation of the *Mor-1* promotor has been associated with the development of chronic post-operative pain (33). Preclinical studies have also highlighted the predisposition to longer post-operative pain in animals exposed to repetitive neonatal procedural pain and, in line with our results, the development of acute pain (22–24). Relevant to the behavioral outcomes is the methylation status of CpG4; the positive correlation observed in our results hints at a methylation-dependent-compensatory mechanism following needle pricks. Moreover, epigenetic regulation of *Mor-1* in the DRG was shown to contribute to reduced morphine analgesia and the development of neuropathic pain (21,28,34,35). Viet and colleagues established that methylation patterns of *Mor-1* are associated with opioid tolerance and pain experience in a mouse model of cancer, a finding that was translated to a clinical population (36). Collectively, the hyper-methylation observed in the *Mor-1* promoter due to repetitive neonatal procedural pain is anticipated to play a role not only in priming the nociceptive system towards prolonged adult post-operative pain but may also increase the susceptibility for neuropathic pain and opioid, e.g. morphine, tolerance.

The present study is the first to establish an impact of repetitive neonatal procedural pain on the methylation of the *Mor-1* and emphasizes the necessity to further explore epigenetic alterations after repetitive stimulation of the developing nociceptive system in order to understand processes driving long-term effects. Preclinical studies have shown that repetitive neonatal procedural pain does not solely affect the nociceptive system, but also has consequences on various cognitive and affective functions and related circuits including the hypothalamus-pituitary-adrenal (HPA) axis (25,37–40). Furthermore, repetitive stimulation of the developing nociceptive system affects neurodevelopment as highlighted by enhanced serotonergic tone in the rostroventral medulla (RVM) of adult rats exposed to repetitive neonatal pain and enhanced firing of dorsal horn spinal cord neurons (7,24). The latter findings are especially relevant in light of recent clinical studies indicating that NICU stay increases the methylation of the serotonin transporter gene (*SLC6A4*), and potentially depletes methylation of Nuclear Receptor Subfamily 3 Group C Member 1 (*NR3C1)* and Solute Carrier Family 1 Member 2 (*SLC1A2)* (18,19,41). These two genes encode a glucocorticoid receptor and a protein involved in glutamate extracellular clearance, respectively (42,43), suggesting that epigenetic alteration following repetitive neonatal procedural pain may have functional implications on neurodevelopment and behavior. Considering the increased *Mor-1* methylation following repetitive needle pricks and evidence from clinical studies, stimulation of the developing nociceptive system likely extends its influence across the epigenome, and thereby yields functional implications still to be understood.

To conclude, repetitive noxious stimulation of the developing nociceptive system has a localized and noxious-stimulus-dependent effect on the methylation of *Mor-1,* in the spinal cord, but not the DRG. Hence, the changes in *Mor-1* methylation status may contribute to the long-lasting behavioral and functional consequences of repetitive neonatal procedural pain.

## Acknowledgments

The completion of this research project would not have been possible without the scientific discussions with colleagues L. Heijmans (Ph.D.), T.J. de Geus (MSc), I. Rudnick-Jansen (Ph.D.), M. Mons (Ph.D.), A. Tiane (Ph.D.), S. Chenine, P. Koulousakis (Ph.D.), and members of the Neuroepigenetics group, Maastricht University, Maastricht, the Netherlands. Optimization of the bisulfite pyrosequencing assays would not have been possible without the help of T. Doeswijk and M. Naldi. We also would like to thank D. Hermes, S. Claessen, and members of the animal facility for their technical support and expertise.

## Authors contribution

All authors substantially contributed to the conception and design of the experiment, interpreted the data and critically revised the manuscript. Acquisition of data and drafting of the article was performed by M. Baudat. The final manuscript was approved by all authors.

## Data availability statement

The data that support the findings of this study are available from the authors upon reasonable request.

## Supplementary materials

**Table S1.**
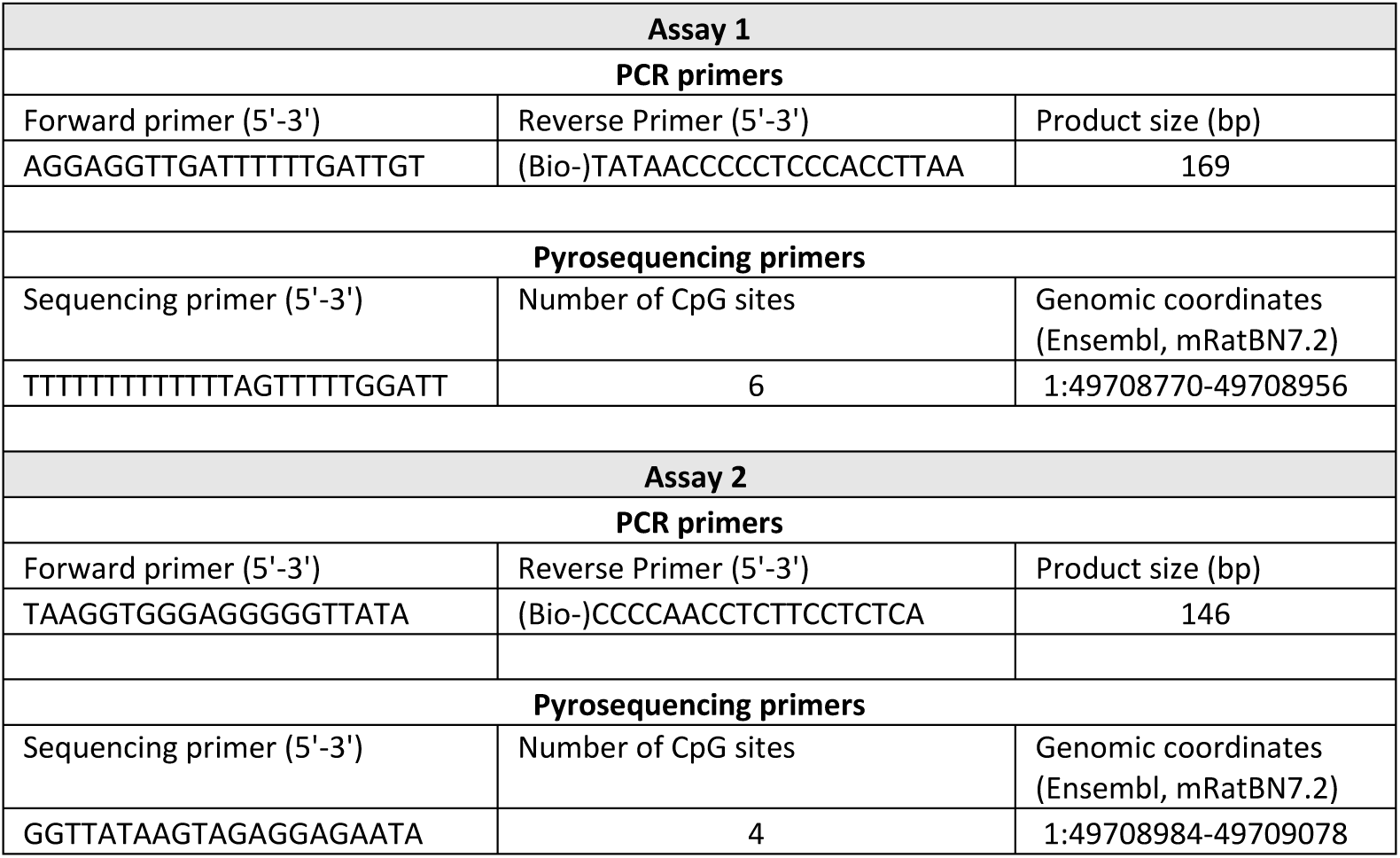
Primers used for bisulfite pyrosequencing of the Mor-1 promotor in the spinal cord and the DRGs of neoantal rats.

**Table S2.** Analysis of spinal cord methylation levels. Spinal cord methylation levels were analyzed with a mix-model ANOVA, and paired t-test were performed when applicable. Only animals that were needle pricked displayed increased methylation levels of the *Mor-1* promotor region on the ipsilateral side compared to the contralateral side. Significant effects are highlighted in bold. Ipsi: Ipsilateral paw; contra: contralateral paw; NP: needle prick (N=10); DC: disturbed control (N=11); UC: undisturbed control (N=11).

| Within-subject variable |  |  |  |  |  |  |  |  | Between-subject variable |
| --- | --- | --- | --- | --- | --- | --- | --- | --- | --- |
|  |  |  |  |  |  | Paired t-tests when applicable |  |  |  |
| Main effect: side | Interaction effect: side*neonatal condition | Main effect: CpG site | Interaction effect: CpG site*neonatal condition | Interaction effect: side*CpG site | Interaction effect: side*CpG site*neonatal condition | NP ipsi. vs NP contra. | DC ipsi. vs DC contra. | UC ipsi. vs UC contra. | Main effect: neonatal condition |
| Promotor methylation levels |  |  |  |  |  |  |  |  |  |
| F(1,28)=5.690, **p=0.024** | F(2,28)=1.02, p=0.374 | - | - | - | - | t(9)=2.683, **p=0.025** | t(10)=1.038, p=0.324 | t(9)=0.565, p=0.586 | F(2,28)=0.559, p=0.578 |
| CpG sites |  |  |  |  |  |  |  |  |  |
| F(1,28)=5.721, **p= 0.024** | F(2,28)=1.004, p=0.379 | F(8,224)=12.949, **p=0.00** | F(16,224)=0.430, p=0.974 | F(8,224)=1.481, p=0.165 | F(16,224)=1.519, p=0.094 | For all CpG sites: p>0.05 | **CpG 4: t(10)=2.925, p=0.015.**<br>For all CpG sites: p>0.05 | For all CpG sites: p>0.05 | F(2,28)=0.224, p=0.801 |

**Table S3.** Analysis of the DRG methylation levels. DRG methylation levels were analyzed with a mix-model ANOVA, and paired t-test were performed when applicable. Neonatal conditions (F2,26=1,965, p=0.106) and sides (ipsilateral vs contralateral) (F1,26=0.709, p=0.408) did not affect methylation levels of the *Mor-1* promotor in the DRGs of P10 animals. DRG: dorsal root ganglion; NP: needle prick (N=10); DC: disturbed control (N=10); UC: undisturbed control (N=10).

| Within-subject variable |  |  |  |  |  | Between-subject variable |
| --- | --- | --- | --- | --- | --- | --- |
| Main effect: side | Interaction effect: side*neonatal condition | Main effect: CpG site | Interaction effect: CpG site*neonatal condition | Interaction effect: side*CpG site | Interaction effect: side*CpG site*neonatal condition | Main effect: neonatal condition |
| **Promotor methylation levels** |  |  |  |  |  |  |
| F(1,26)=0.709, p=0.408 | F(2,26)=0.292, p=0.749 | - | - | - | - | F(2,26)=1.965, p=0.106 |
| **CpG sites** |  |  |  |  |  |  |
| F(1,26)=0.696, p=0.412 | F(2,26)=0.116, p=0.891 | F(8,208)=62.827, **p=0.00** | F(16,208)=0.613, p=0.872 | F(8,208)=0.387, p=0.927 | F(16,208)=0.703, p=0.789 | F(2,27)=1.029, p=0.205 |

**Table S4.** Correlation of behavioral data with methylation of the Mor-1 in the spinal cord. Correlation of the last PWT measured, that is at P7 five hours after the last needle prick or disturbtion, with the average spinal methylation levels of *Mor −1* and CpG site specific methylation levels. Rs: Spearman correlation coefficient. NP: Needle pricks, N= 10. DC: Distrubed control, N=11.

|  | PWT,P7+5h | *Mor-1* promotor methylation | CpG1 | CpG2 | CpG3 | CpG4 | CpG5 | CpG6 | CpG7 | CpG8 | CpG9 |
| --- | --- | --- | --- | --- | --- | --- | --- | --- | --- | --- | --- |
| NP | Ipsilateral | $R_s=0.152$, $p=0.676$ | $R_s=0.612$, $p=0.060$ | $R_s=-0.588$, $p=0.074$ | $R_s=-0.043$, $p=0.907$ | **$R_s=0.855$, $p=0.002$** | $R_s=0.164$, $p=0.651$ | $R_s=-0.018$, $p=0.960$ | $R_s=0.109$, $p=0.763$ | $R_s=0.576$, $p=0.082$ | $R_s=0.188$, $p=0.603$ |
| | Contralateral | $R_s=0.222$, $p=0.537$ | $R_s=0.068$, $p=0.852$ | $R_s=0.401$, $p=0.250$ | $R_s=0.554$, $p=0.096$ | $R_s=0.043$, $p=0.906$ | $R_s=-0.445$, $p=0.198$ | $R_s=0.599$, $p=0.067$ | $R_s=-0.143$, $p=0.693$ | $R_s=0.136$, $p=0.708$ | $R_s=-0.435$, $p=0.209$ |
| DC | Ipsilateral | $R_s=-0.046$, $p=0.893$ | $R_s=0.475$, $p=0.140$ | $R_s=-0.240$, $p=0.478$ | $R_s=-0.155$, $p=0.650$ | $R_s=-0.358$, $p=0.280$ | $R_s=0.138$, $p=0.685$ | $R_s=0.143$, $p=0.675$ | $R_s=0.035$, $p=0.919$ | $R_s=-0.143$, $p=0.675$ | $R_s=0.300$, $p=0.371$ |
| | Contralateral | $R_s=0.447$, $p=0.168$ | $R_s=-0.205$, $p=0.544$ | $R_s=0.251$, $p=0.456$ | $R_s=0.379$, $p=0.250$ | $R_s=-0.237$, $p=0.482$ | $R_s=0.449$, $p=0.166$ | $R_s=-0.199$, $p=0.557$ | $R_s=0.132$, $p=0.698$ | $R_s=0.338$, $p=0.309$ | $R_s=0.307$, $p=0.359$ |

## References

1. Blencowe H, Cousens S, Chou D, et al. Born too soon: the global epidemiology of 15 million preterm births. Reprod Health 2013;10:1–14.

2. Tielsch JM. Global incidence of preterm birth. In: Low-Birthweight Baby: Born Too Soon or Too Small. Karger Publishers; 2015. p. 9–15.

3. Gamber RA, Blonsky H, Mcdowell M, Lakshminrusimha S. Declining birth rates, increasing maternal age and neonatal intensive care unit admissions. Journal of Perinatology [Internet] 2024 [cited 2024 Mar 21];44:203–8. Available from: 10.1038/s41372-023-01834-x

4. Carbajal R, Rousset A, Danan C, et al. Epidemiology and treatment of painful procedures in neonates in intensive care units. JAMA 2008;300:60–70.

5. Cruz MD, Fernandes AM, Oliveira CR. Epidemiology of painful procedures performed in neonates: a systematic review of observational studies. European Journal of Pain 2016;20:489–98.

6. Grunau RE. Neonatal pain in very preterm infants: long-term effects on brain, neurodevelopment and pain reactivity. Rambam Maimonides Med J 2013;4.

7. de Kort AR, Joosten EA, Patijn J, Tibboel D, van den Hoogen NJ. Anatomical changes in descending serotonergic projections from the rostral ventromedial medulla to the spinal dorsal horn following repetitive neonatal painful procedures. International Journal of Developmental Neuroscience 2022;82:361–71.

8. van den Hoogen NJ, de Kort AR, Allegaert KM, et al. Developmental neurobiology as a guide for pharmacological management of pain in neonates. Semin Fetal Neonatal Med. 2019;24.

9. LaPrairie JL, Murphy AZ. Neonatal Injury Alters Adult Pain Sensitivity by Increasing Opioid Tone in the Periaqueductal Gray. Front Behav Neurosci [Internet] 2009 [cited 2023 Feb 16];3. Available from: https://www.readcube.com/articles/10.3 389%2Fneuro.08.031.2009

10. Kwok CHT, Devonshire IM, Bennett AJ, Hathway GJ. Postnatal maturation of endogenous opioid systems within the periaqueductal grey and spinal dorsal horn of the rat. Pain 2014;155:168–78.

11. Hathway GJ, Vega-Avelaira D, Fitzgerald M. A critical period in the supraspinal control of pain: opioid-dependent changes in brainstem rostroventral medulla function in preadolescence. Pain 2012;153:775–83.

12. Arvidsson U, Fliedi M, Chakrabarti S, et al. Distribution and Targeting of a p.-Opioid Receptor (MORI) in Brain and Spinal Cord. The Journal of Neuroscience 1995;75:3328–41.

13. Terman GW, Eastman CL, Chavkin C. Mu Opiates Inhibit Long-Term Potentiation Induction in the Spinal Cord Slice. 2001 [cited 2024 Feb 28];Available from: www.jn.physiology.org

14. van den Hoogen NJ, van Reij RRI, Patijn J, Tibboel D, Joosten EAJ. Adult spinal opioid receptor μ1 expression after incision is altered by early life repetitive tactile and noxious procedures in rats. Dev Neurobiol 2018;78:417–26.

15. Klose RJ, Bird AP. Genomic DNA methylation: the mark and its mediators. TRENDS in Biochemical Sciences [Internet] 2006 [cited 2024 Jan 22];31:89–97. Available from: www.sciencedirect.com

16. Jirtle RL, Skinner MK. Environmental epigenomics and disease susceptibility. Nat Rev Genet. 2007;8:253–62.

17. Kundakovic M, Jaric I. The epigenetic link between prenatal adverse environments and neurodevelopmental disorders. Genes (Basel). 2017;8.

18. Van Dokkum NH, Bao M, Nynke Verkaik-Schakel R, et al. Neonatal stress exposure and DNA methylation of stress-related and neurodevelopmentally relevant genes: An exploratory study. Early Hum Dev [Internet] 2023 [cited 2024 Jan 25];186:105868. Available from: http://creativecommons.org/licenses/by/4.0/

19. Provenzi L, Fumagalli M, Sirgiovanni I, et al. Pain-related stress during the neonatal intensive care unit stay and SLC6A4 methylation in very preterm infants. Front Behav Neurosci 2015;9.

20. Chau CMY, Ranger M, Sulistyoningrum D, Devlin AM, Obertander TF, Grunau RE. Neonatal pain and comt Val158Met genotype in relation to serotonin transporter (SLC6A4) promoter methylation in very preterm children at school age. Front Behav Neurosci 2014;8.

21. Sun N, Yu L, Gao Y, et al. MeCP2 Epigenetic Silencing of Oprm1 Gene in Primary Sensory Neurons Under Neuropathic Pain Conditions. Front Neurosci 2021;15.

22. Knaepen L, Patijn J, van Kleef M, Mulder M, Tibboel D, Joosten EAJ. Neonatal repetitive needle pricking: plasticity of the spinal nociceptive circuit and extended postoperative pain in later life. Dev Neurobiol 2013;73:85–97.

23. van den Hoogen NJ, Patijn J, Tibboel D, Joosten EA. Repetitive noxious stimuli during early development affect acute and long-term mechanical sensitivity in rats. Pediatr Res 2020;87:26–31.

24. van den Hoogen NJ, Patijn J, Tibboel D, Joosten BA, Fitzgerald M, Kwok CHT. Repeated touch and needle-prick stimulation in the neonatal period increases the baseline mechanical sensitivity and postinjury hypersensitivity of adult spinal sensory neurons. Pain 2018;159:1166–75.

25. de Kort AR, Joosten EA, Patijn J, Tibboel D, van den Hoogen NJ. Neonatal procedural pain affects state, but not trait anxiety behavior in adult rats. Dev Psychobiol 2021;63:e22210.

26. Heinke B, Gingl E, Sandkühler J. Cellular/Molecular Multiple Targets of-Opioid Receptor-Mediated Presynaptic Inhibition at Primary Afferent A-and C-Fibers. 2011 [cited 2024 Feb 28];Available from: http://rsb.info.nih.gov/ij/

27. Hatfield LA, Hoffman RK, Polomano RC, Conley Y. Epigenetic Modifications Following Noxious Stimuli in Infants. Biol Res Nurs 2018;20:137–44.

28. Sun L, Zhao J-Y, Gu X, et al. Nerve injury–induced epigenetic silencing of opioid receptors controlled by DNMT3a in primary afferent neurons. Pain 2017;158:1153–65.

29. Hwang CK, Song KY, Kim CS, et al. Evidence of Endogenous Mu Opioid Receptor Regulation by Epigenetic Control of the Promoters. Mol Cell Biol [Internet] 2007 [cited 2024 Feb 23];27:4720–36. Available from: https://www.tandfonline.com/action/journalInformation?journalCode=tmcb20

30. Beggs S, Currie G, Salter MW, Fitzgerald M, Walker SM. Priming of adult pain responses by neonatal pain experience: maintenance by central neuroimmune activity. Brain 2012;135:404–17.

31. Walker SM, O’reilly H, Beckmann J, Marlow N. Conditioned pain modulation identifies altered sensitivity in extremely preterm young adult males and females. 2018;

32. Walker SM, Melbourne A, O’Reilly H, et al. Somatosensory function and pain in extremely preterm young adults from the UK EPICure cohort: sex-dependent differences and impact of neonatal surgery. Br J Anaesth 2018;121:623–35.

33. Chidambaran V, Zhang X, Martin LJ, et al. Dna methylation at the mu-1 opioid receptor gene (OPRM1) promoter predicts preoperative, acute, and chronic postsurgical pain after spine fusion. Pharmgenomics Pers Med [Internet] 2017 [cited 2024 Feb 23];10:157–68. Available from: https://www.tandfonline.com/action/journalInformation?journalCode=dpgp20

34. Mo K, Wu S, Gu X, et al. MBD1 Contributes to the Genesis of Acute Pain and Neuropathic Pain by Epigenetic Silencing of Oprm1 and Kcna2 Genes in Primary Sensory Neurons. Journal of Neuroscience [Internet] 2018 [cited 2024 Feb 28];38:9883–99. Available from: 10.1523/JNEUROSCI.08 80-18.2018

35. Zhou X-L, Yu L-N, Wang Y, et al. Increased methylation of the MOR gene proximal promoter in primary sensory neurons plays a crucial role in the decreased analgesic effect of opioids in neuropathic pain. 2012 [cited 2024 Feb 23];Available from: http://www.molecularpain.com/content/10/1/51

36. Viet CT, Dang D, Aouizerat BE, et al. OPRM1 Methylation Contributes to Opioid Tolerance in Cancer Patients. Journal of Pain 2017;18:1046–59.

37. Davis SM, Rice M, Rudlong J, Eaton V, King T, Burman MA. Neonatal pain and stress disrupts later-life pavlovian fear conditioning and sensory function in rats: Evidence for a two-hit model. Dev Psychobiol 2018;60:520–33.

38. Mooney-leber SM, Brummelte S. Neonatal pain and reduced maternal care alter adult behavior and hypothalamic– pituitary–adrenal axis reactivity in a sex specific manner. Dev Psychobiol 2020;62:631–43.

39. Chen M, Xia D, Min C, et al. Neonatal repetitive pain in rats leads to impaired spatial learning and dysregulated hypothalamic-pituitary-adrenal axis function in later life. Sci Rep 2016;6:1–14.

40. Baudat M, Simons SHP, Joosten EAJ. Repetitive neonatal procedural pain affects stress-induced plasma corticosterone increase in young adult females but not in male rats. Dev Psychobiol [Internet] 2024 [cited 2024 Mar 4];Available from: https://onlinelibrary.wiley.com/doi/10.1002/dev.22478

41. Oberlander TF, Weinberg J, Papsdorf M, Grunau R, Misri S, Devlin AM. Prenatal exposure to maternal depression, neonatal methylation of human glucocorticoid receptor gene (NR3C1) and infant cortisol stress responses. Epigenetics 2008;3:97–106.

42. Desilva TM, Kinney HC, Borenstein NS, et al. The Glutamate Transporter EAAT2 Is Transiently Expressed in Developing Human Cerebral White Matter. J Comp Neurol [Internet] 2007 [cited 2024 Mar 4];501:879–90. Available from: https://onlinelibrary.wiley.com/doi/10.1002/cne.21289

43. Palma-Gudiel H, Córdova-Palomera A, Leza JC, Fañanás L. Glucocorticoid receptor gene (NR3C1) methylation processes as mediators of early adversity in stress-related disorders causality: A critical review. Neurosci Biobehav Rev. 2015;55:520–35.

